# *Hierarchical* Clustering on RNA Dependent RNA Polymerase using Machine Learning

**DOI:** 10.1101/2021.08.23.457366

**Authors:** Gudipati Pavan Kumar

**Affiliations:** Independent Researcher, Hyderabad, India

**Keywords:** RNA Dependent RNA Polymerase, RdRP, Machine Learning, *Hierarchical* Clustering, sklearn, scipy, TfidfVectorizer, Dendrogram, Biopython

## Abstract

RNA Dependent RNA Polymerase (RdRP) catalyzes the replication of RNA from an RNA template and is mostly found in Viruses. We have collected over 161 viral RdRP FASTA Sequences from the NCBI protein database using python script. Each of these sequences was transformed with TfidfVectorizer using sklearn module, with the “one Letter” word, because each Letter belongs to one Amino acid. These transformed data were sent to Hierarchical clustering using scipy library and visualized data using Dendrogram. These Machine Learning technique is able to classify or segment similar RdRp into one cluster. Each of these clusters was tested for their multiple sequence alignment with COBALT of NCBI. We observed that these clusters predicted similar RdRP among various viruses. These techniques can be further improved to segment or classify various proteins. These Machine Learning or Artificial Intelligence techniques need more improvement in their algorithms to solve genomics and proteomics.

**Repository:** Programs and Datasets : https://github.com/DSPavan/RdRP

## I. INTRODUCTION

**RNA-dependent RNA polymerase** (**RdRp**) is an enzyme that catalyzes the replication of RNA from an RNA template. RNA dependent RNA polymerase (RdRp) of SARS COV-2 is known to mediate replication of the viral genome and its propagation inside host cells [1]. Several RNA containing Viruses have this enzyme. ***Hierarchical clustering***, also known as *hierarchical cluster analysis*, is an algorithm that groups similar objects into groups called *clusters*. The endpoint is a set of clusters, where each cluster is distinct from each other cluster, and the objects within each cluster are broadly similar to each other. [2,3]. NCBI Protein database is a collection of sequences from several sources, including translations from annotated coding regions in GenBank, RefSeq and TPA, as well as records from SwissProt, PIR, PRF, and PDB. [4]. Python is a programing language widely used in Machine Learning, and they have ***sklearn*** [5] and ***scipy*** [6] modules. Main aim of this experiment is to test, can we cluster similar RdRP with help of Machine Learning / Artificial Intelligence techniques?

## II. MATERIAL & METHODS

### A) Collection of RdRP

Primarily this data collected from NCBI website, under Protein Database. These data can be collected manually from this database, using search option. We can also Biopython modules to get all these data from programmatically. I have extracted RdRP data using Biopython [7] with Entrez [8] module, specifically Amino acid sequences length between 600 to 3500 and excluded partial, probable, putative types from the list, so that we will get clean records. I have used kaggle notebook [9] with Python, which have preinstalled biopython libraries. I have collected 490+ RdRP from various organisms, mostly Viruses. After extraction of data, manually curation is done, to remove duplicates or identical proteins. After final curation we have dataset of 161 RdRP proteins.

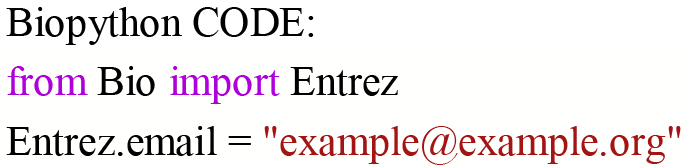

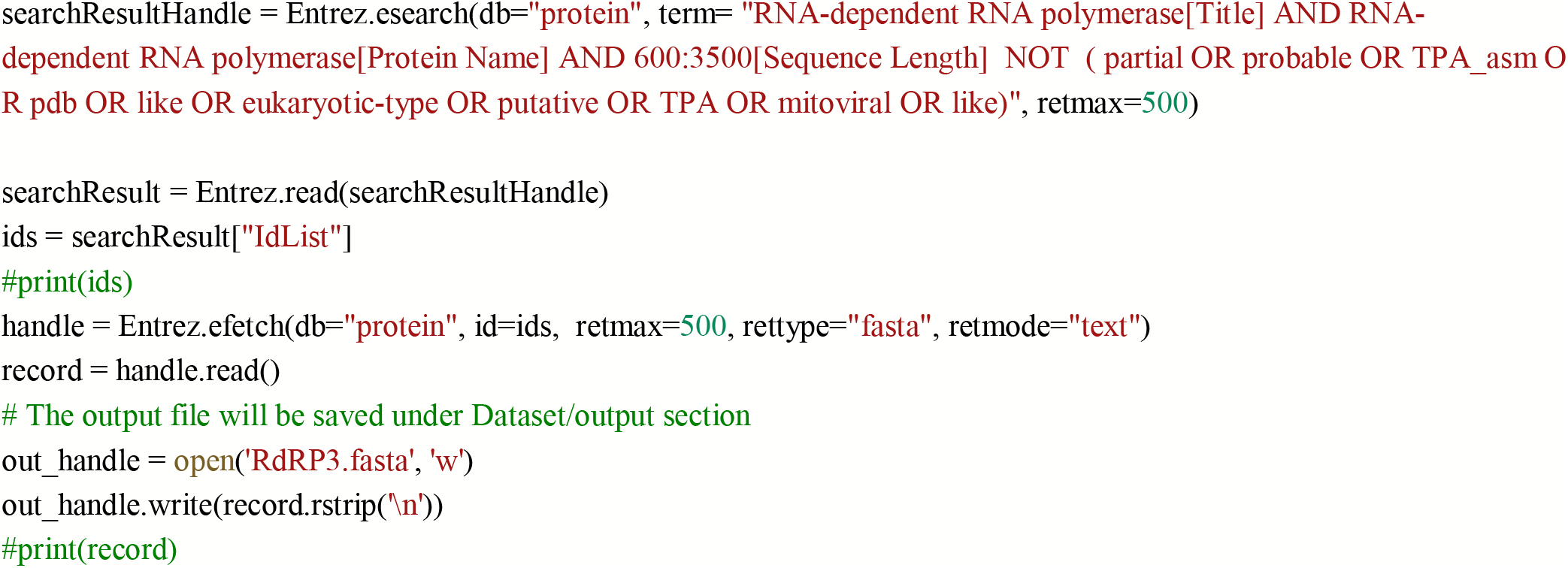

### B) TfidfVectorizer

Each Fasta sequences of RdRP was transformed with TfidfVectorizer method with in sklearn module. TfidfVectorizer, Convert a collection of raw documents to a matrix of TF-IDF features. In this experiment, we specifically tried with single letter word, so that, each amino acid in Fasta sequence can be transformed properly with above matrix.

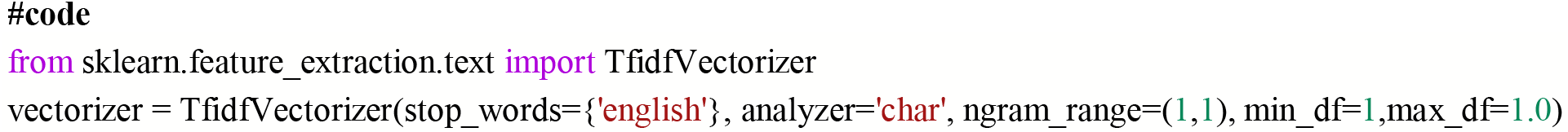

### C) Using Kmeans

Number of optimum clusters were identified with Elbow technique and silhouette coefficients. Optimum Cluster to get all 161 Fasta sequences was 10 to 20 clusters.

### D) Hierarchical Cluster

**After** TfidfVectorizer, this data clustered with Linkage type “ward”. Then it is converted into Dendogram tree. After verifying dendrogram results, we can observe 8 major clusters, with color_threshold=0.14, each of these can be given unqiue color, so that, we can analyze each cluster for further analysis. You can vary color threshold, so that, you can verify different cluster sizes. I have kept this value (0.14), so that we can optimum size of clusters for analysis. With Cluster size-15, all 161 sequences clearly assigned to specific cluster.

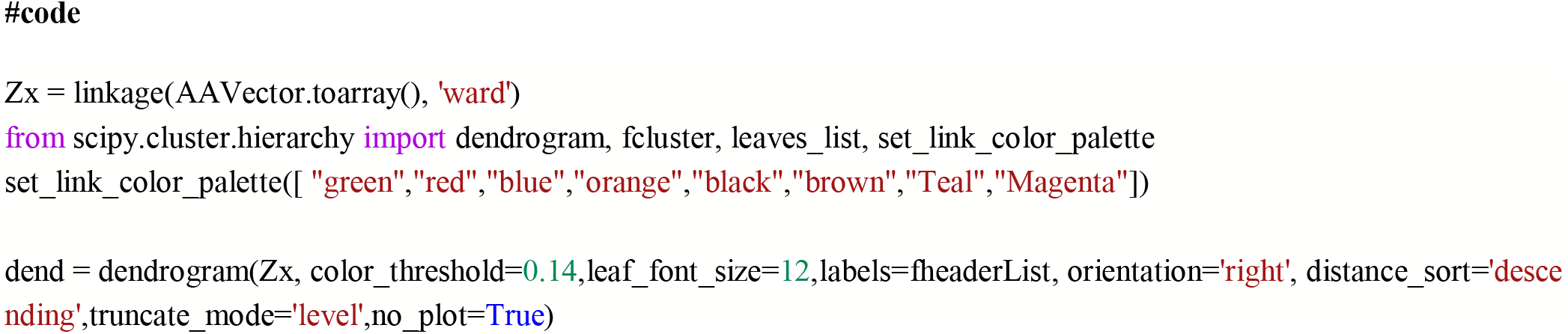

### E) Extraction of Each Cluster for Further Analysis

We will get complete Denogram from above code, We need to extract, each color Cluster, using fcluster module, and we can extract list of fasta sequences of RdRP, which are grouped in same cluster.

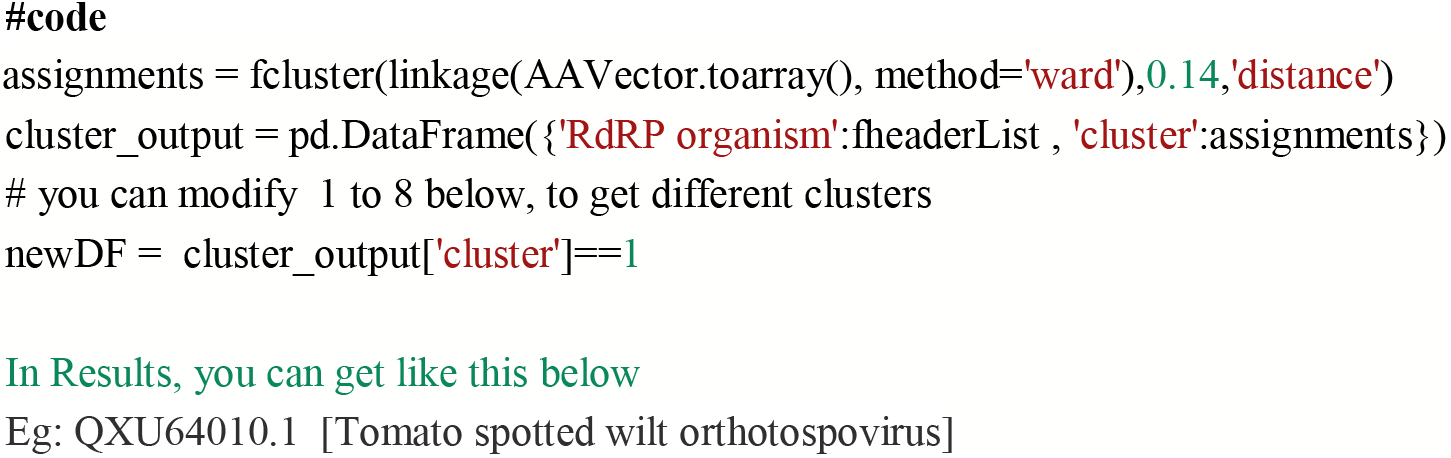

### F) Constraint-based Multiple Alignment Tool of NCBI

Each of these clusters, which we extracted from above, was given as input (Accession Number is sufficient) into COBALT (Constraint-based Multiple Alignment Tool of NCBI) [10, 11]. After alignment, set Conservation Setting as “Identity”. We collected, near to cluster branch as in dendrogram tree results, and send again for alignment, and we got below results.

## III. RESULTS AND DISCUSSION

Each of these clusters sequences are aligned with COBALT-(Constraint-based Multiple Alignment Tool) with Conservation Setting as Identity.

### A. Green Cluster

**Fig. 1:**
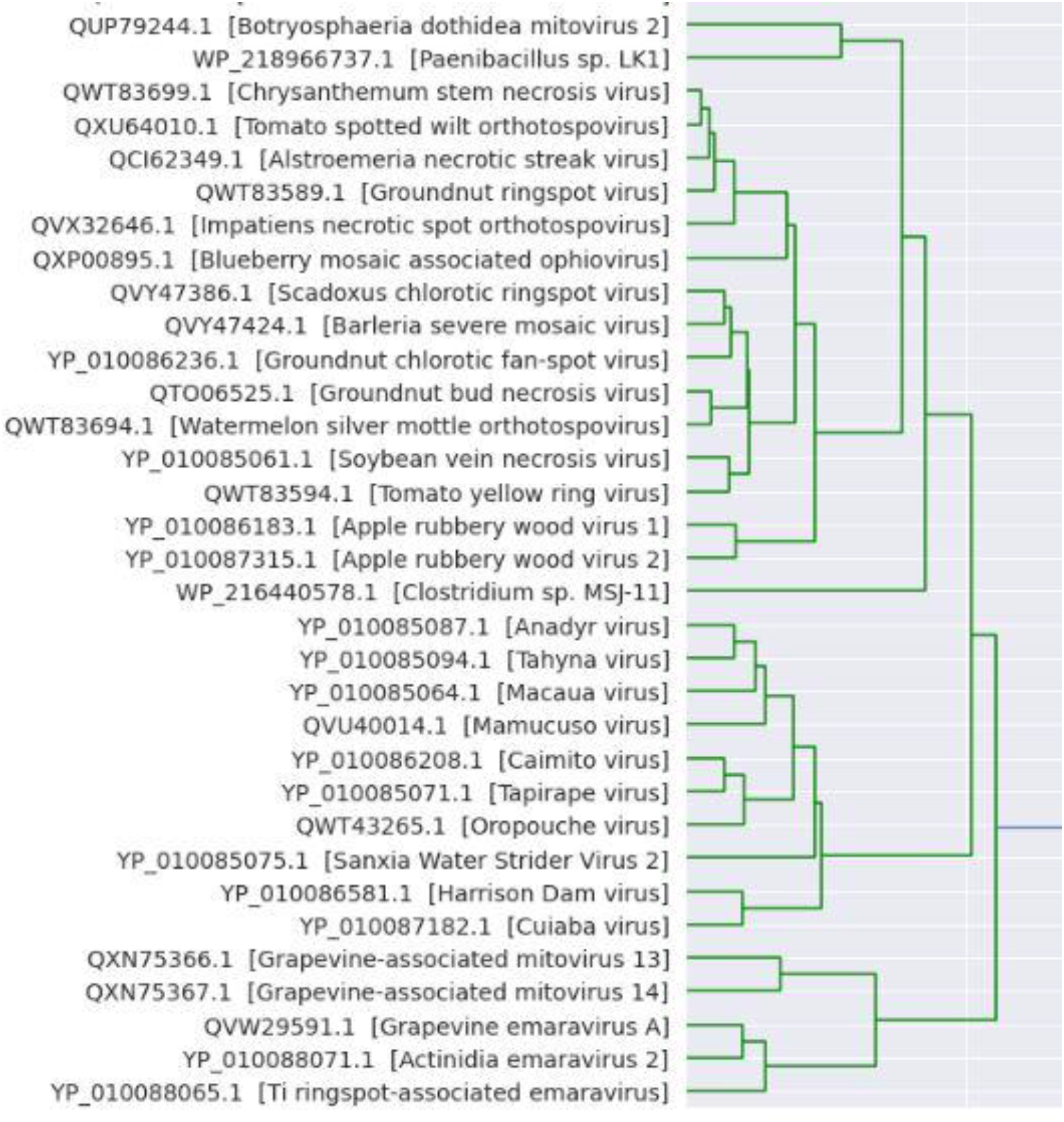
Green Cluster.

**Table 1:**
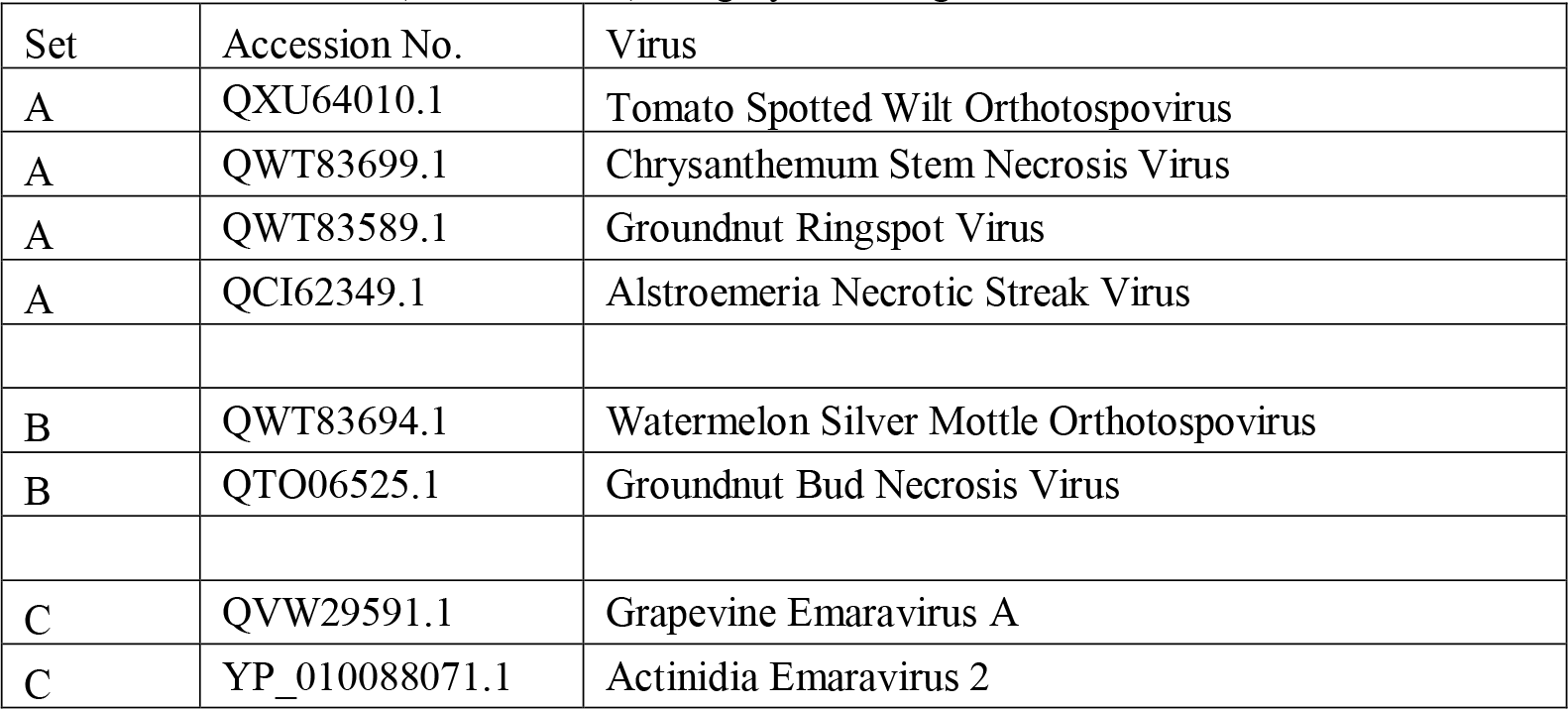
Green Cluster (Cluster No: 1) : Highly matching cluster

### B. Red Cluster

**Fig. 2:**
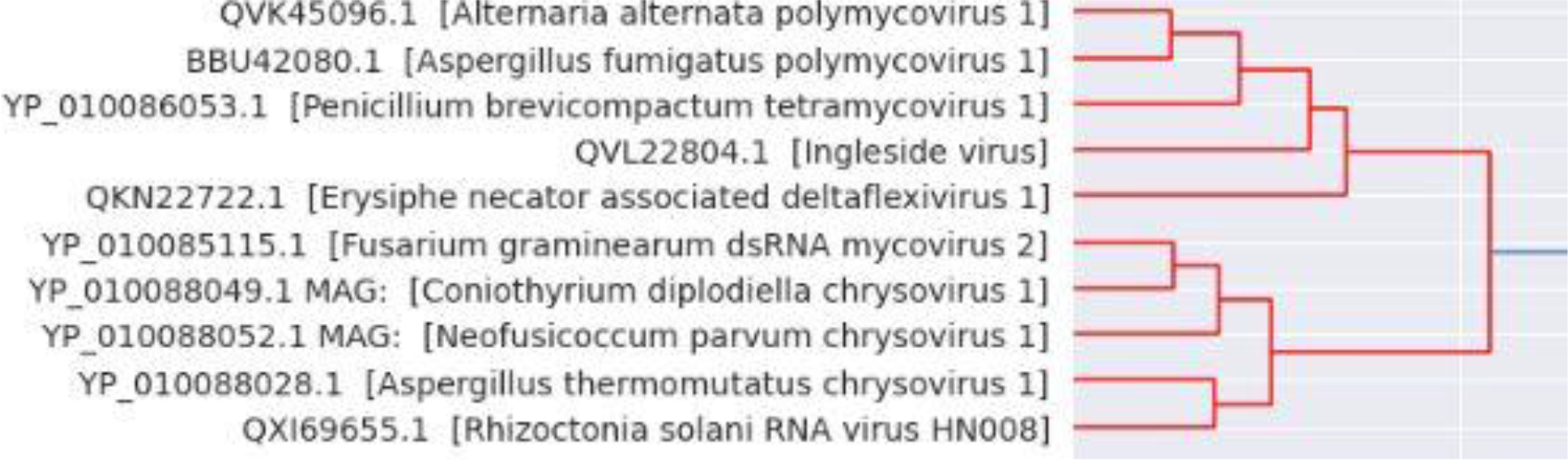
Red Cluster.

**Table 2:**
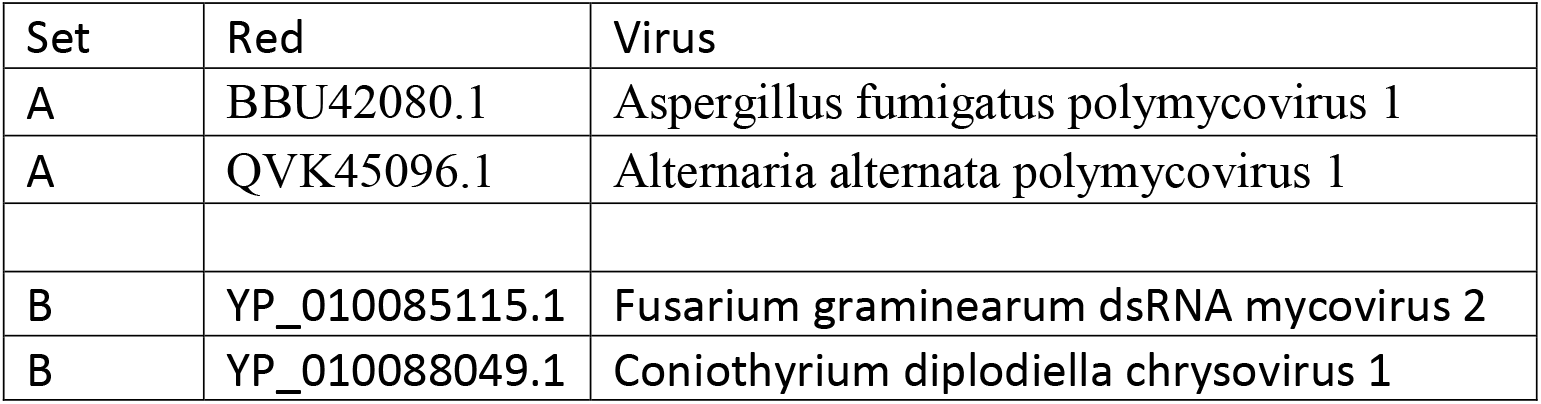
Red Cluster (Cluster No: 2): Highly matching cluster

### C. Blue Cluster

**Table 3:**
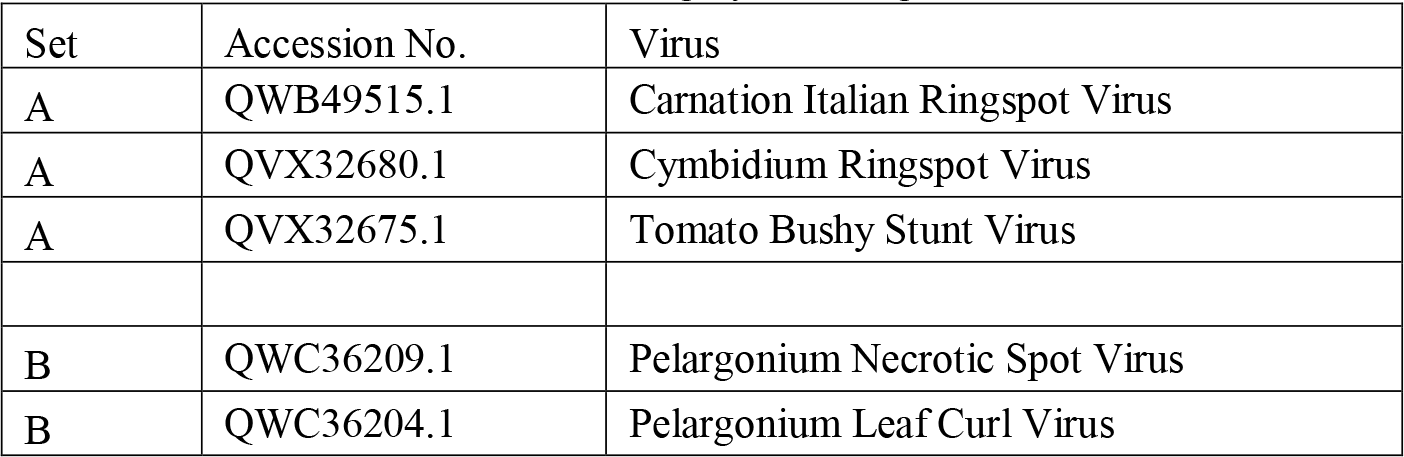
Blue Cluster (Cluster No: 3) Highly matching cluster

**Fig. 3:**
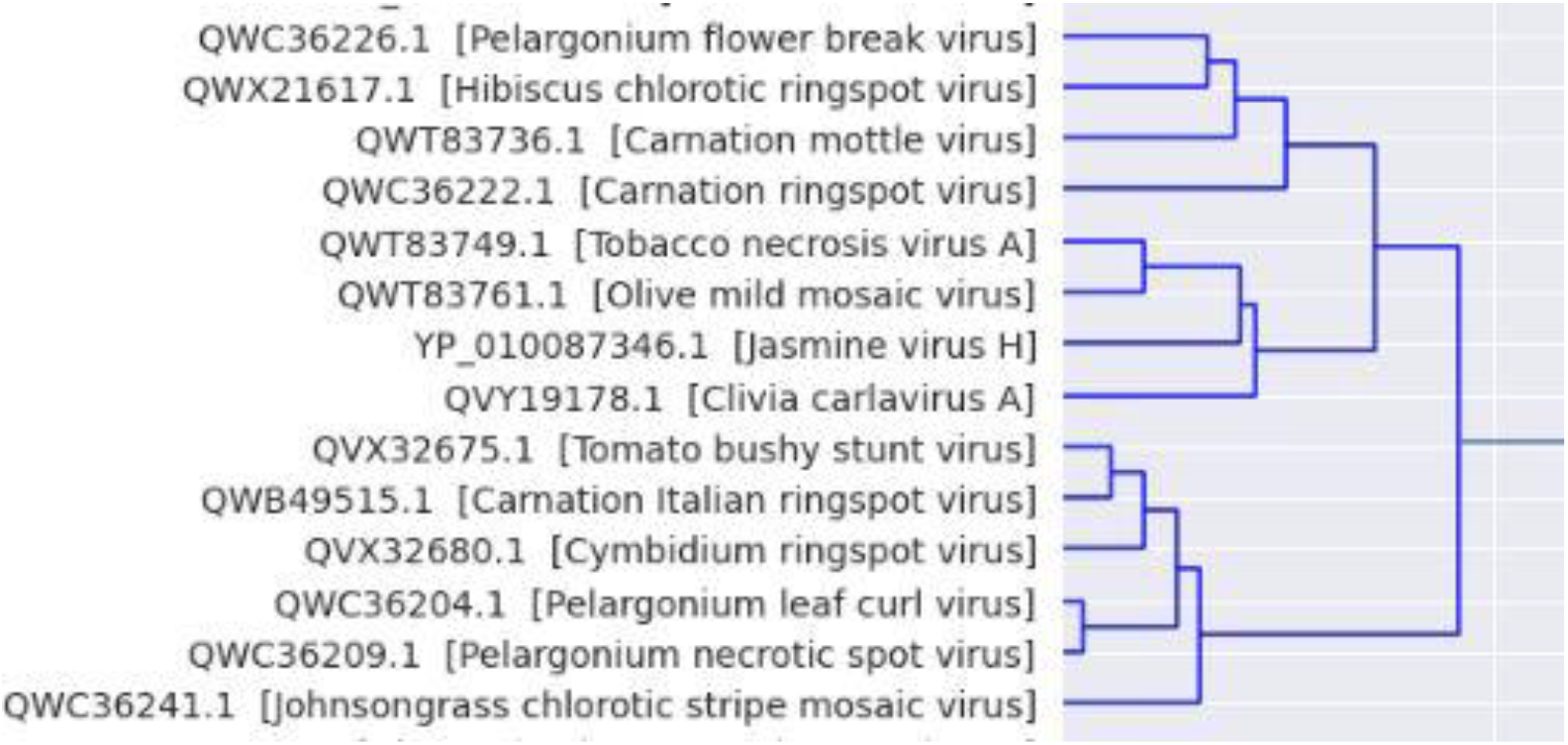
Blue Cluster.

### D. Orange Cluster

**Fig. 4:**
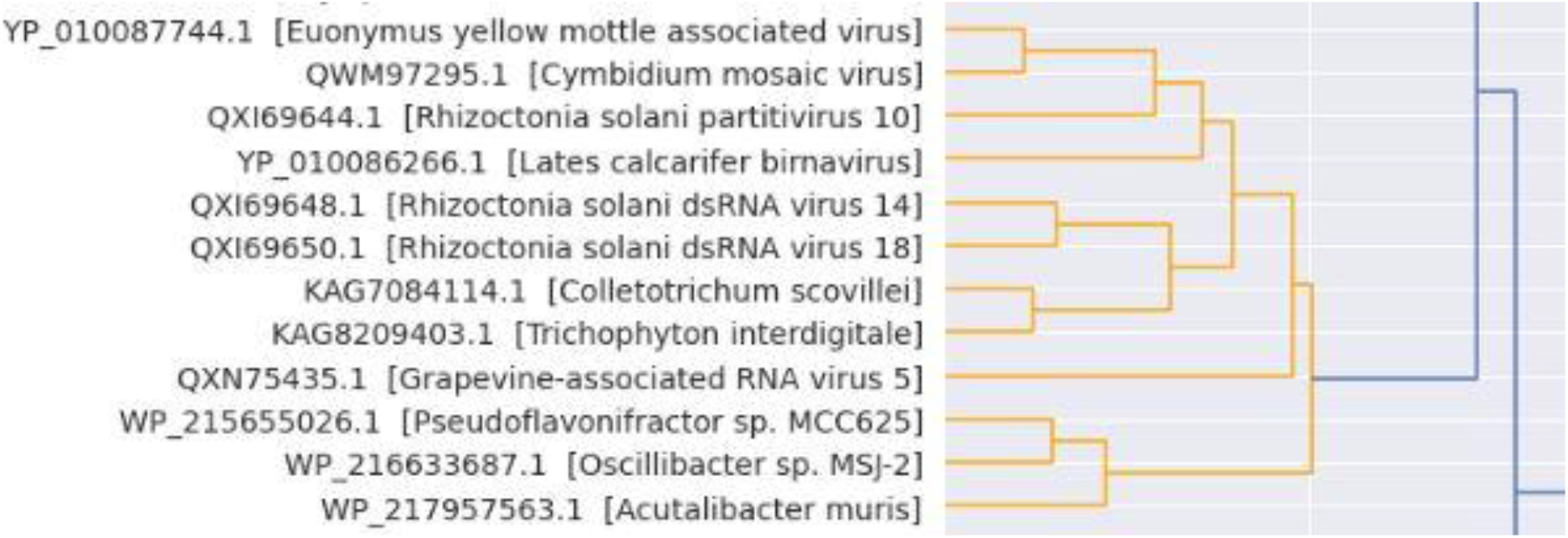
Orange Cluster.

**Table 4:**
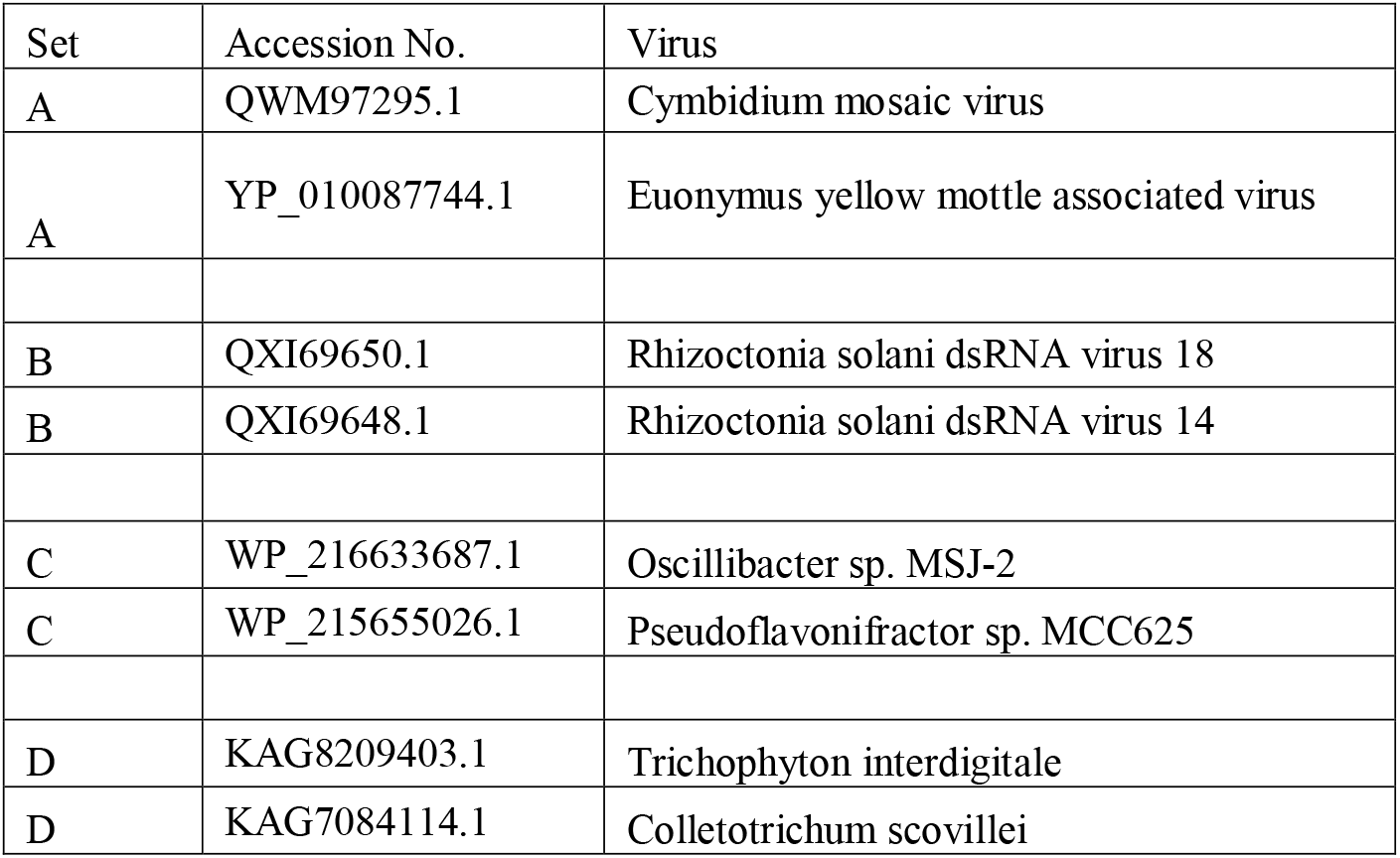
Orange Cluster (Cluster No: 4): Highly matching cluster

### E. Black Cluster

**Fig. 5:**
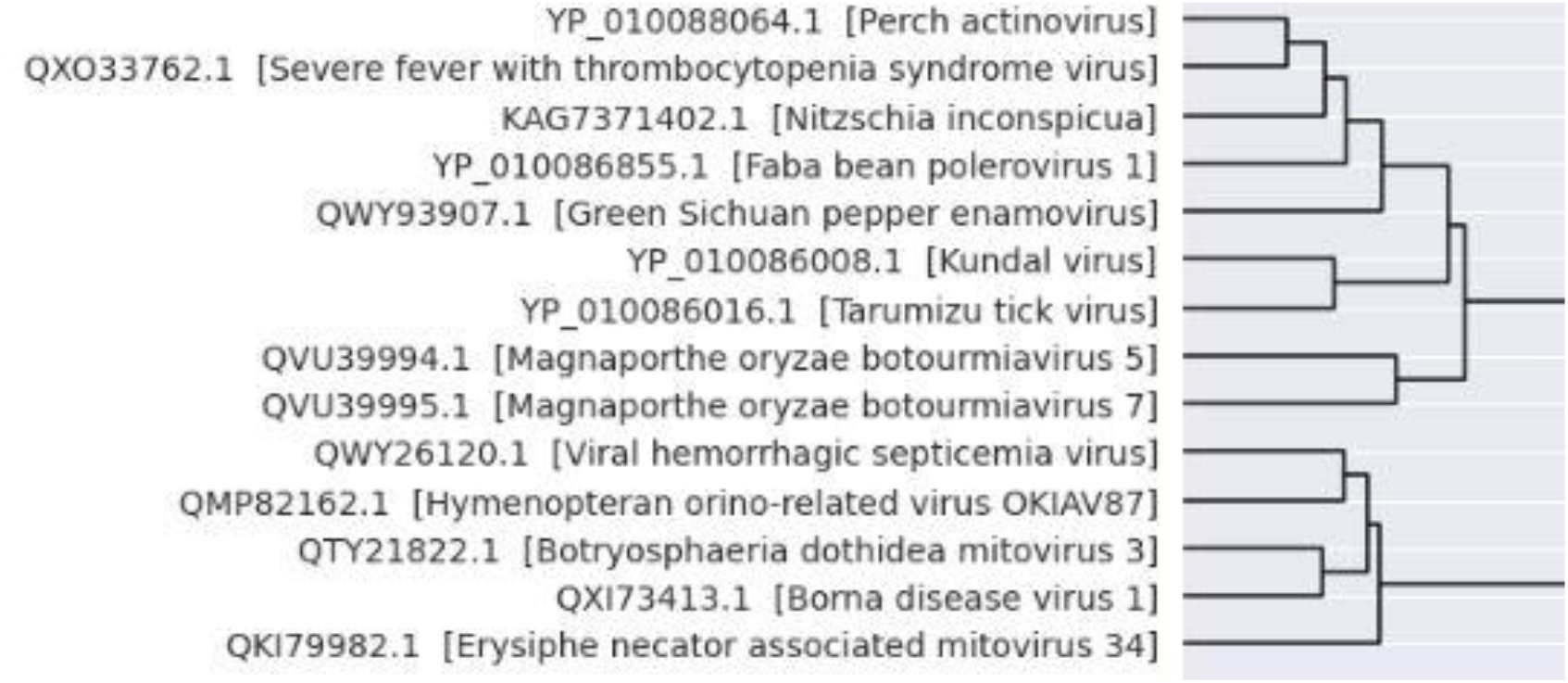
Black Cluster.

**Table 5:**
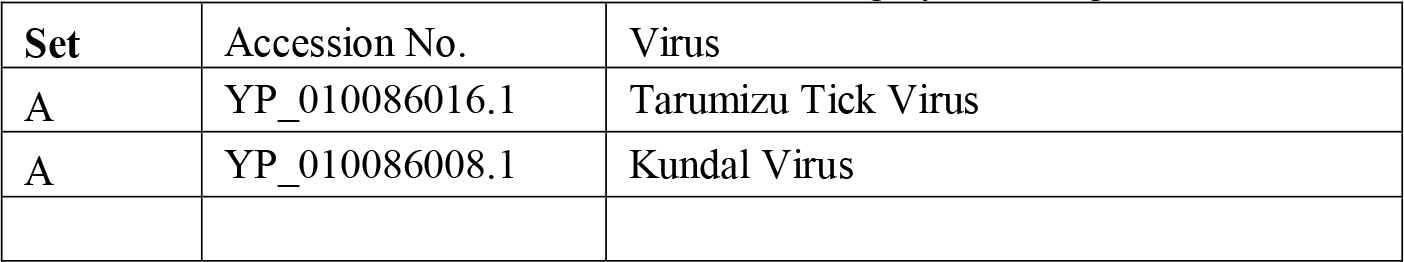
Black Cluster (Cluster No: 5) : Highly matching cluster **Note**: There is no good match with other sequences, and huge difference is observed, probably because of long distance between them.

| Set | Accession No. | Virus |
| --- | --- | --- |
| A | YP_010086016.1 | Tarumizu Tick Virus |
| A | YP_010086008.1 | Kundal Virus |

### F. Brown Cluster

**Table 6:**
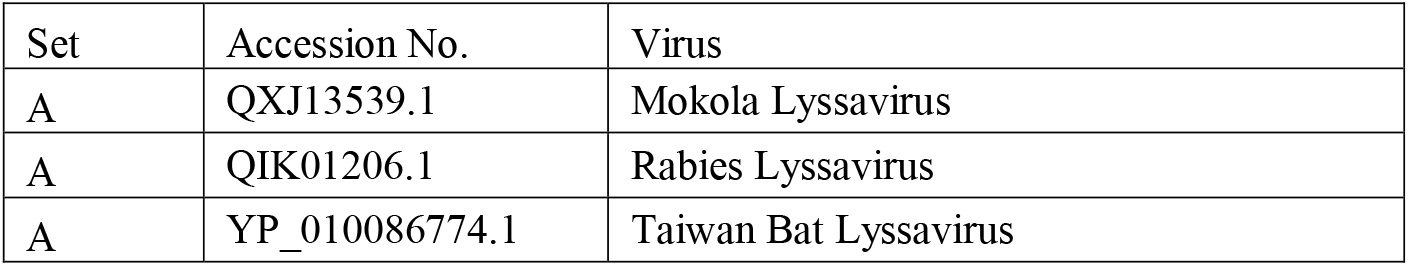
Brown Cluster (Cluster No: 6) : Highly matching cluster

| Set | Accession No. | Virus |
| --- | --- | --- |
| A | QXJ13539.1 | Mokola Lyssavirus |
| A | QIK01206.1 | Rabies Lyssavirus |
| A | YP_010086774.1 | Taiwan Bat Lyssavirus |

**Fig. 6:**
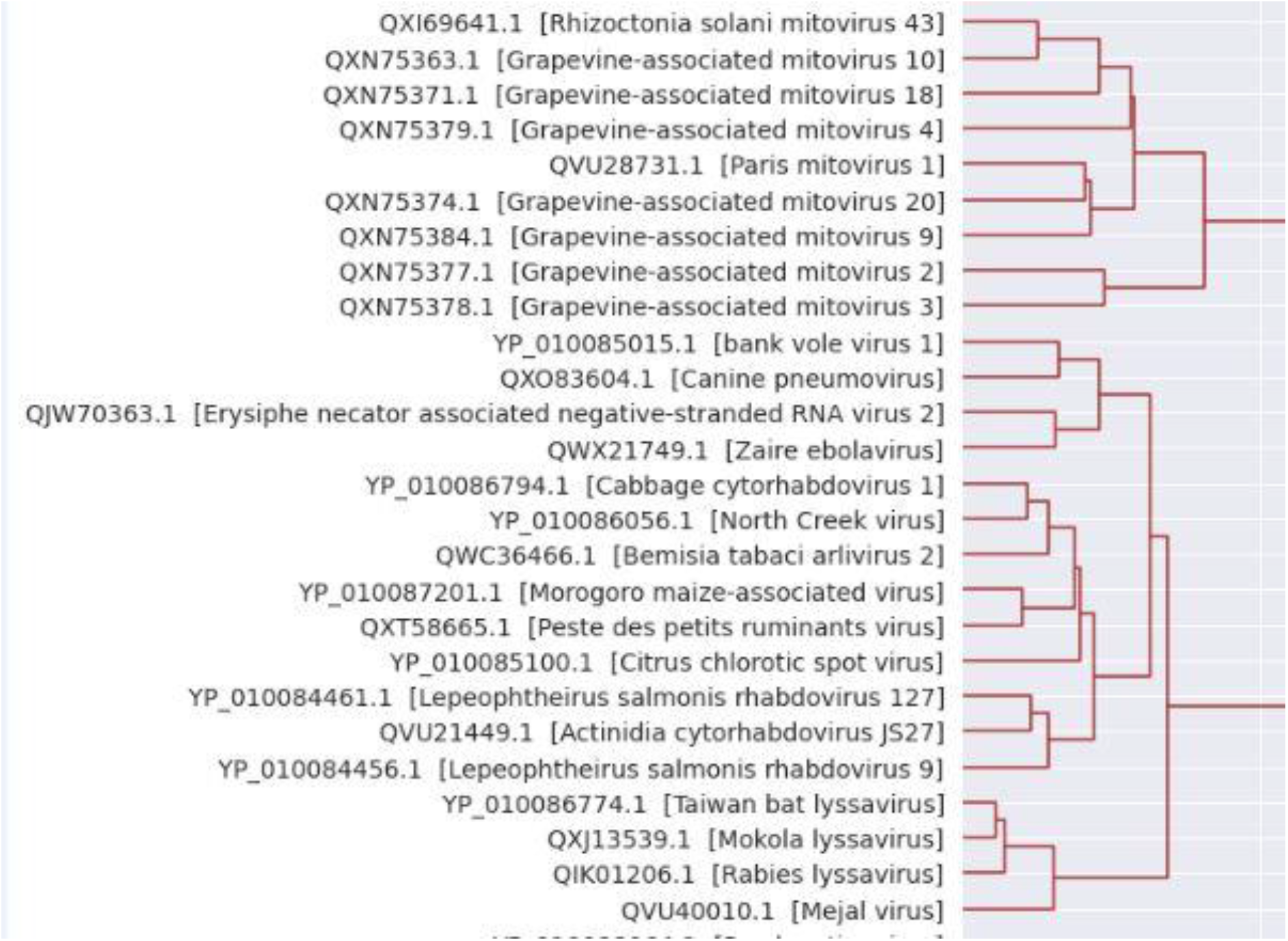
Brown Cluster.

### G. Teal Cluster

**Fig. 7:**
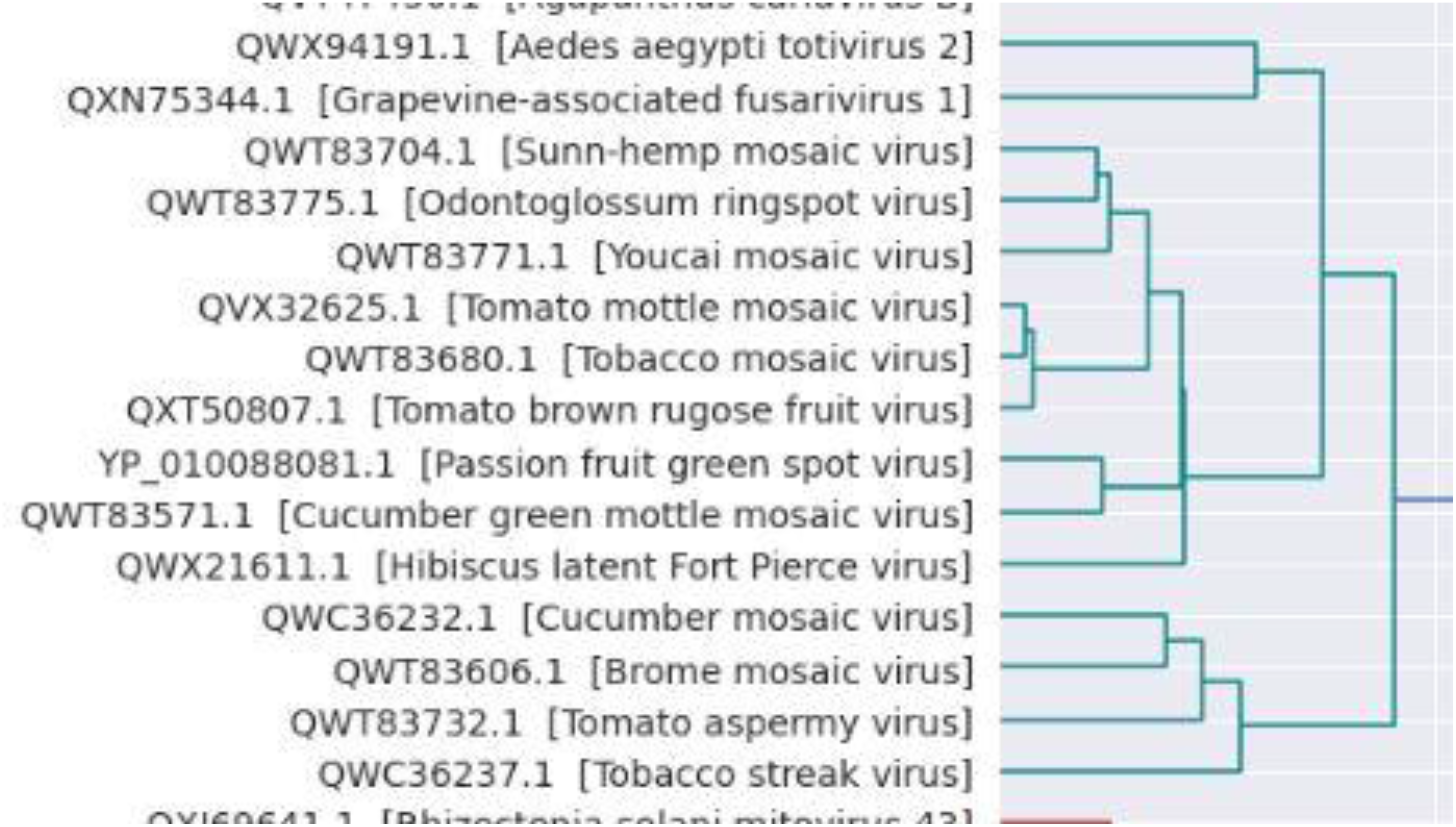
Teal Cluster.

**Table 7:**
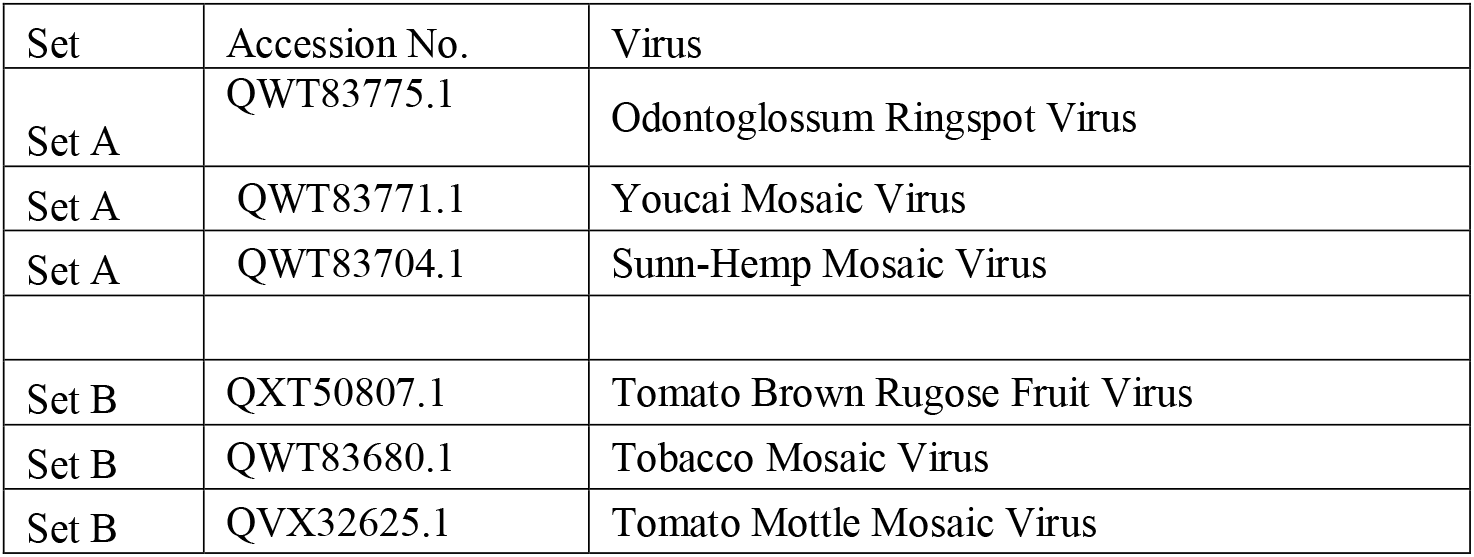
Teal Cluster (Cluster No: 7) : Highly matching cluster

### H. Magneta Cluster

**Fig. 8:**
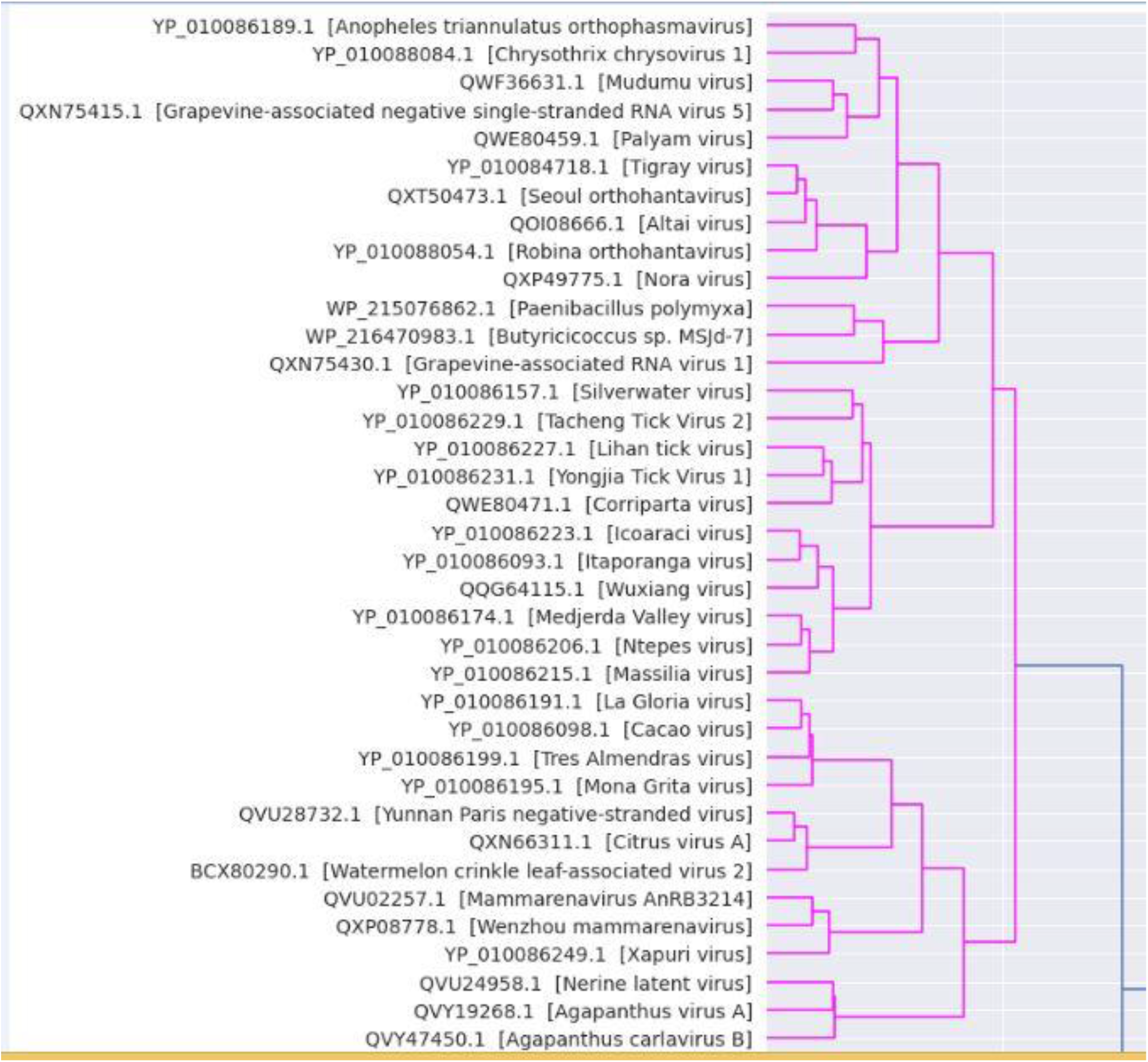
Magneta Cluster.

**Table 8:**
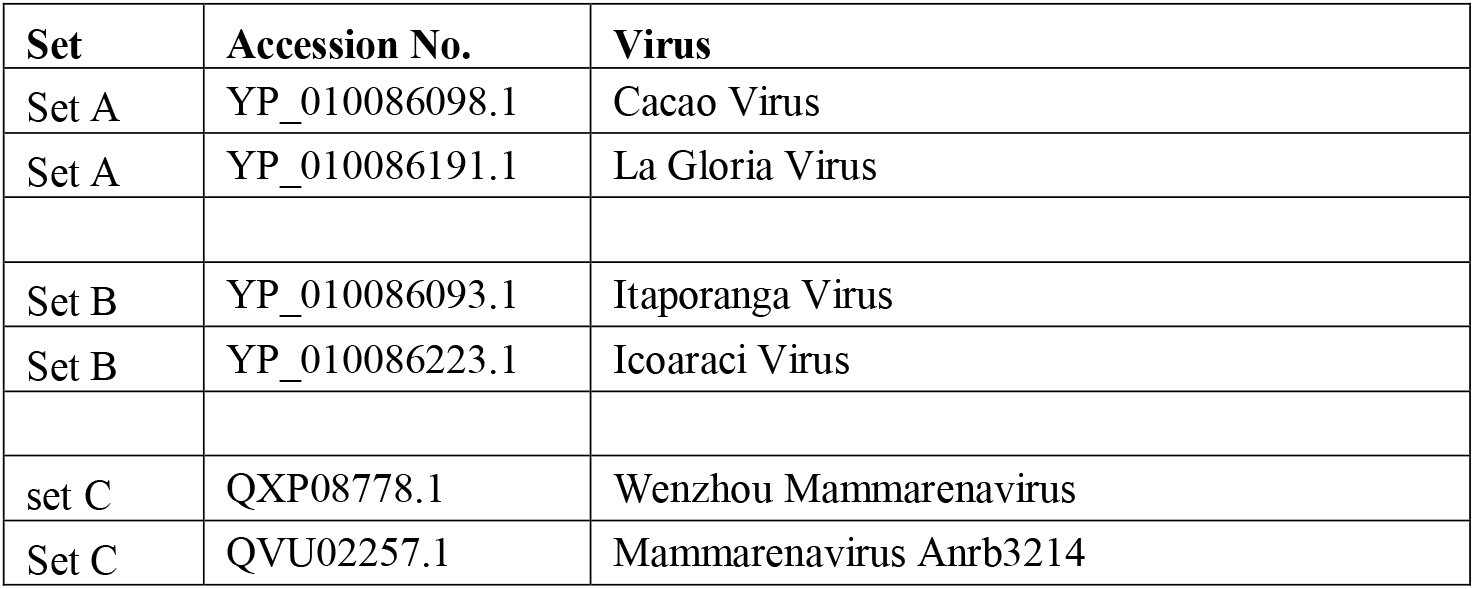
Magneta Cluster (Cluster No: 8): Highly matching cluster

**Fig. 9:**
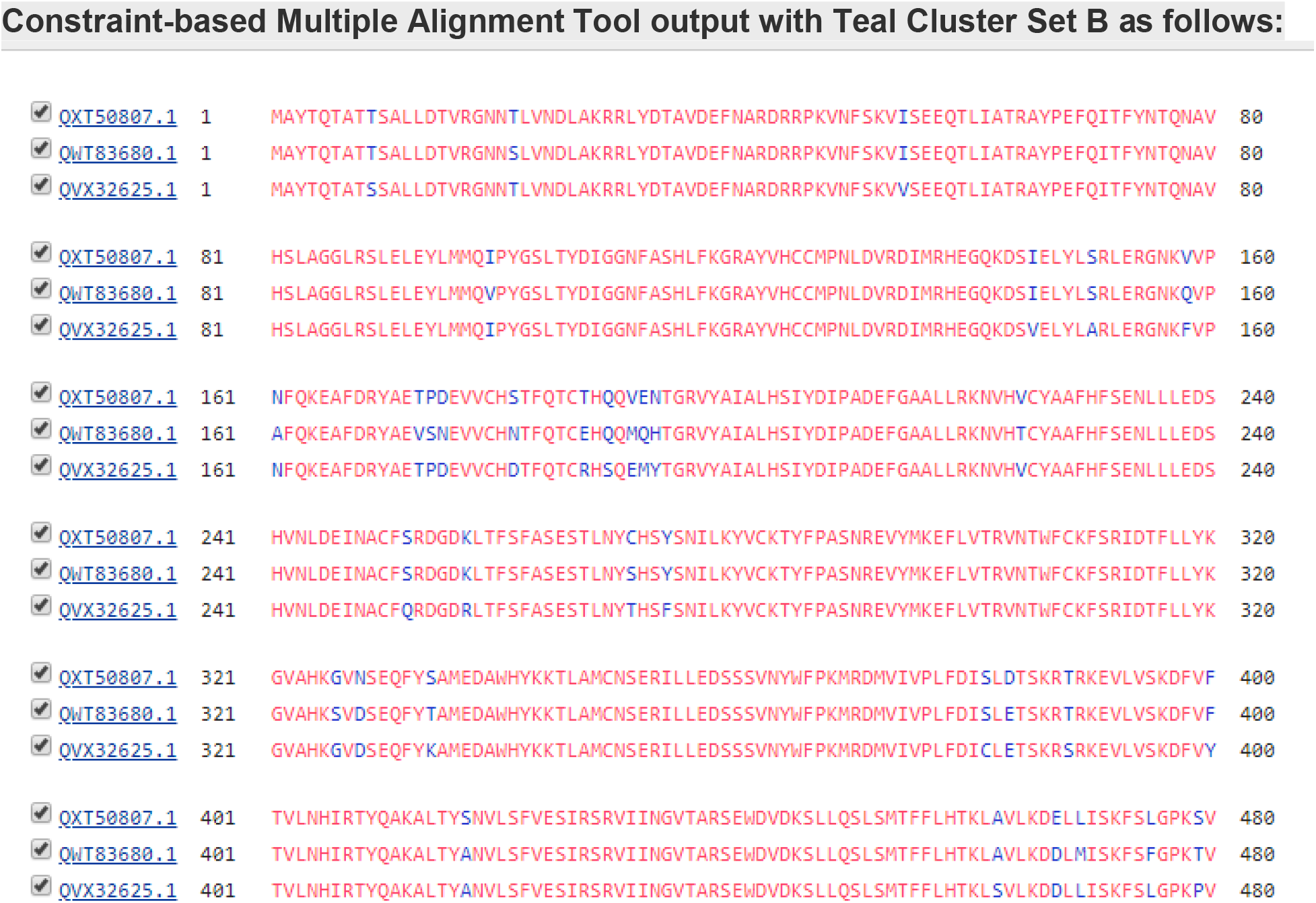
COBALT output for Teal Cluster Set B.

## IV. CONCLUSIONS

Hierachical clustering technique is able to identify similar groups based on Protein sequence. We able to identify similar groups eg.1: Tomato spotted wilt orthotospovirus, Chrysanthemum stem necrosis virus, Groundnut ringspot virus, Alstroemeria necrotic streak virus another example Rhizoctonia solani RNA virus HN008, Aspergillus fumigatus polymycovirus 1, Erysiphe necator associated deltaflexivirus 1, Alternaria alternata polymycovirus 1, Ingleside virus. All these possible due to Machine Learning techniques such as Hierarchical clustering method. These Methods or algorithms can be further improved to solve genomics and proteomics problems.

## Supporting information

cobalt results

## VII. ACKNOWLEDGMENT

This paper and the research behind it would not have been possible without Python, Biopython, NCBI and Friend’s support.

---

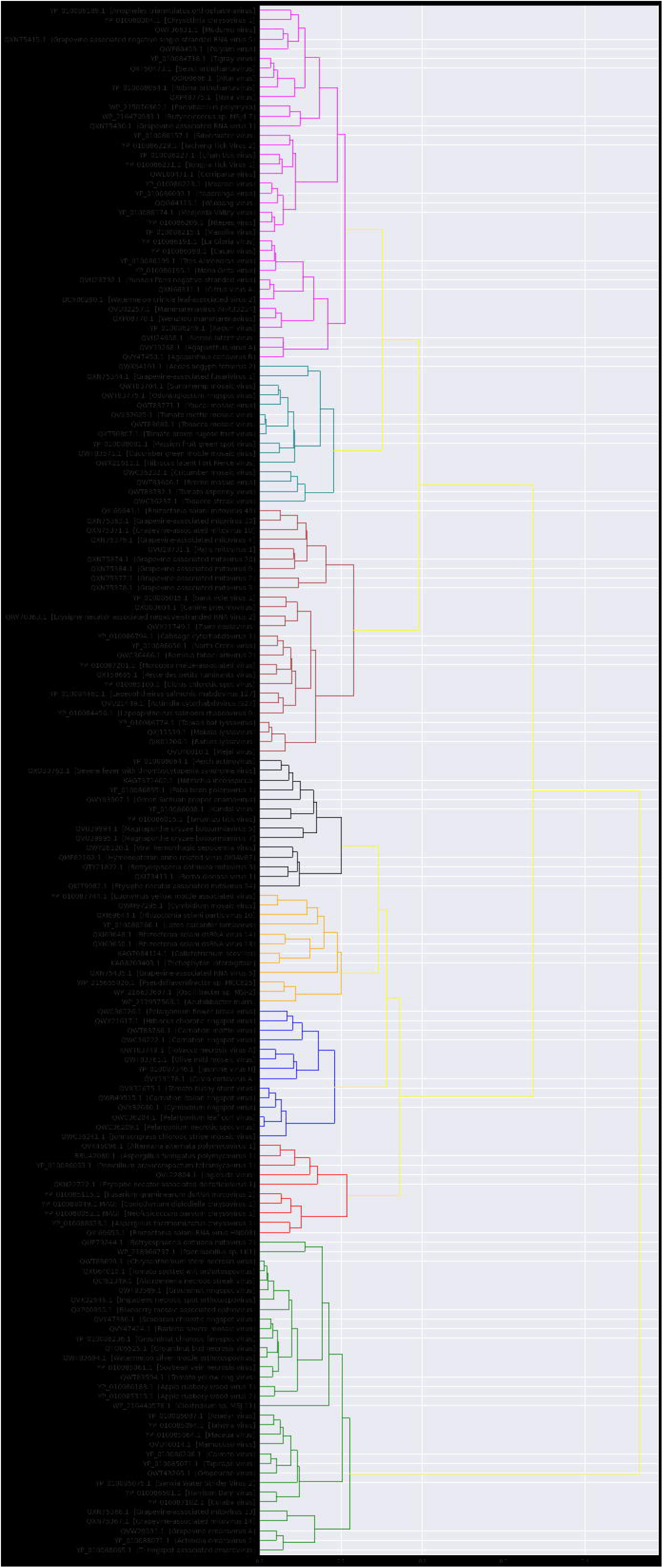

