## Supplementary material for "*Hierarchical* Clustering on RNA Dependent RNA Polymerase using Machine Learning": cobalt results

A)
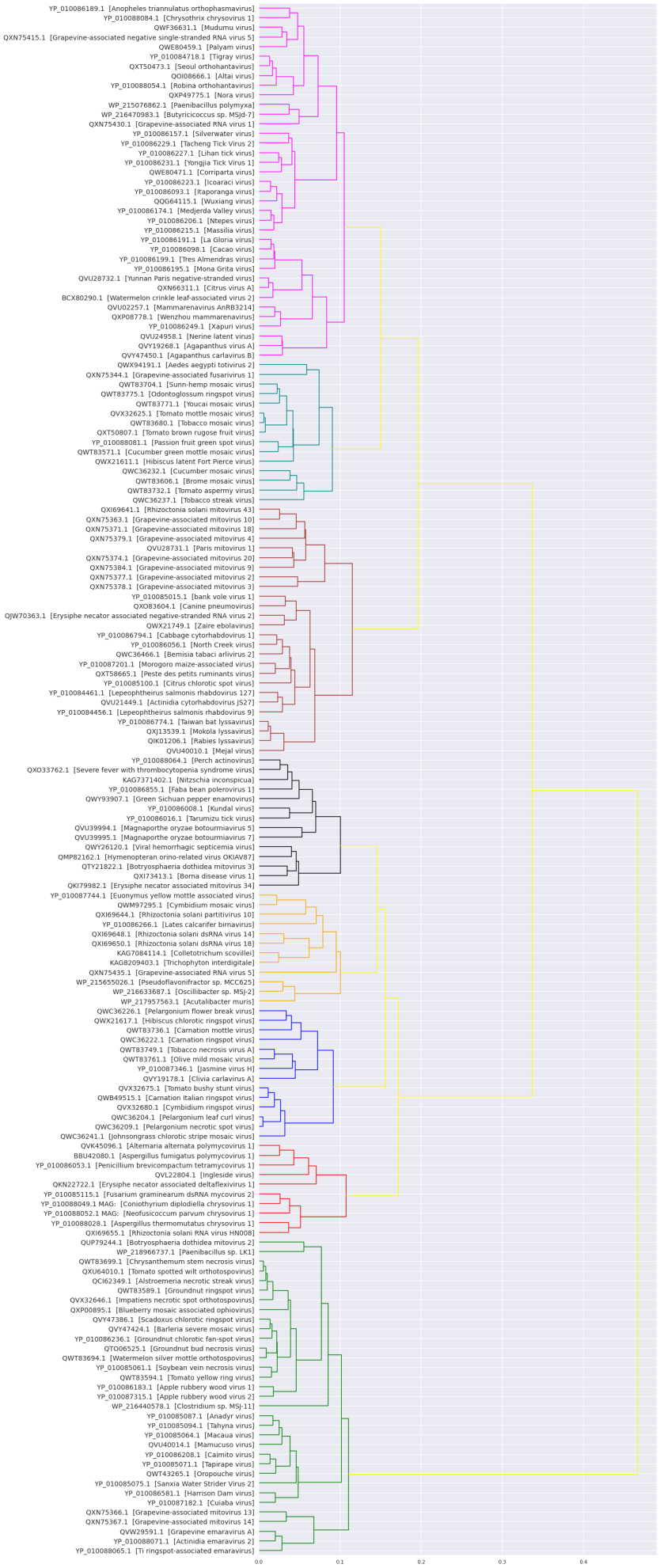
Result:

Complete Dendrogram

B ) Cluster Outputs

(1). Green Cluster

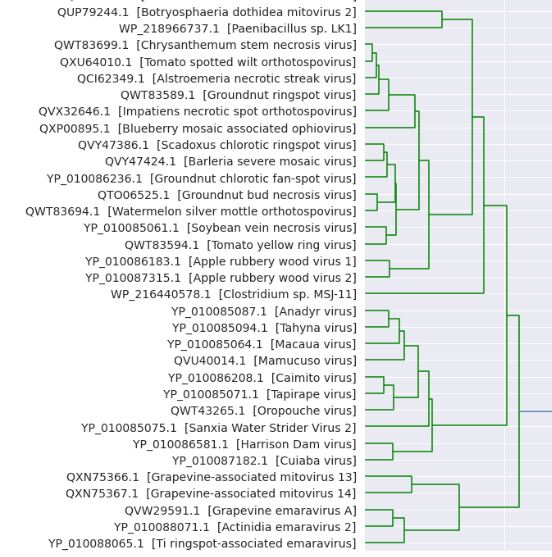

2) Red Color :

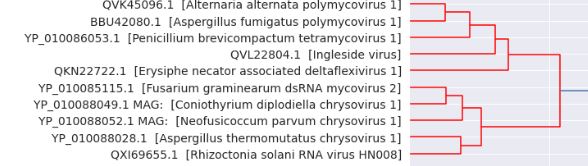

3) Blue:

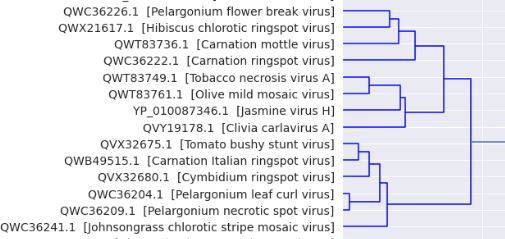

4) Orange:

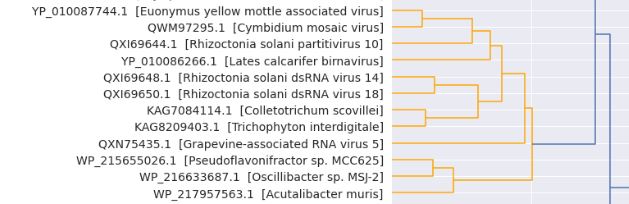

5) Black

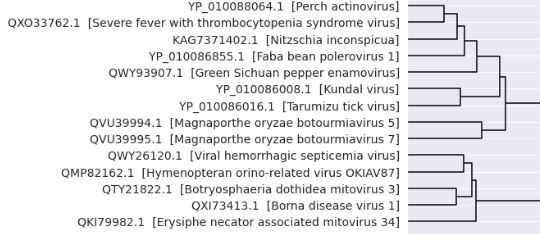

6) Brown:

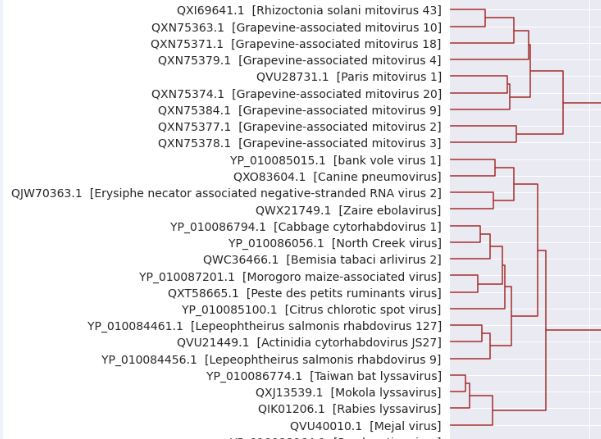

7) Teal

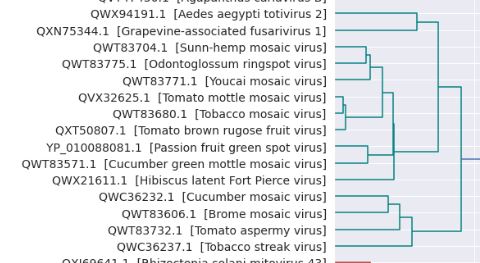

8) Magneta

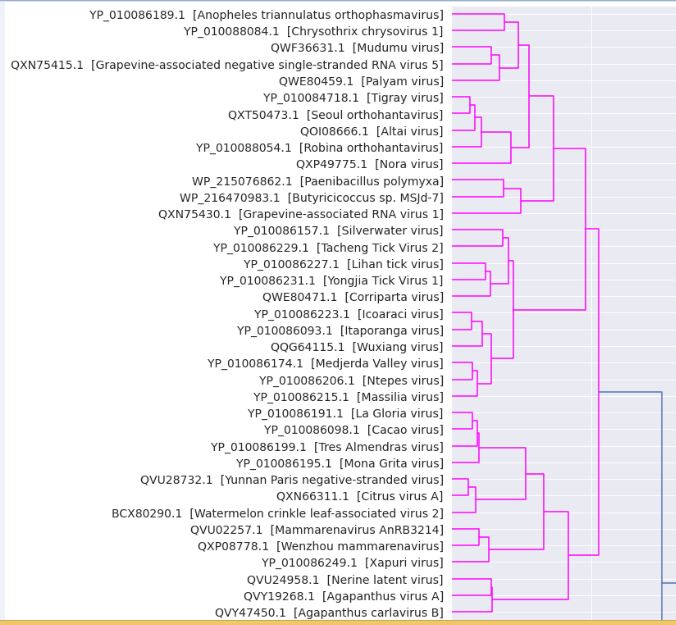

C) Highly conserved matching in clusters.

Green Cluster (Cluster No: 1)

| Set | Green | Virus |
| --- | --- | --- |
| A | QXU64010.1 | Tomato spotted wilt orthotospovirus |
| A | QWT83699.1 | Chrysanthemum stem necrosis virus |
| A | QWT83589.1 | Groundnut ringspot virus |
| A | QCI62349.1 | Alstroemeria necrotic streak virus |
| B | QWT83694.1 | Watermelon silver mottle orthotospovirus |
| B | QTO06525.1 | Groundnut bud necrosis virus |
| C | QVW29591.1 | Grapevine emaravirus A |
| C | YP_010088071.1 | Actinidia emaravirus 2 |

Red Cluster (Cluster No: 2)

| Set | Red | Virus |
| --- | --- | --- |
| A | BBU42080.1 | Aspergillus fumigatus polymycovirus 1 |
| A | QVK45096.1 | Alternaria alternata polymycovirus 1 |
| B | YP_010085115.1 | Fusarium graminearum dsRNA mycovirus 2 |
| B | YP_010088049.1 | Coniothyrium diplodiella chrysovirus 1 |

Blue Cluster (Cluster No: 3)

| Set | Blue | Virus |
| --- | --- | --- |
| A | QWB49515.1 | Carnation Italian ringspot virus |
| A | QVX32680.1 | Cymbidium ringspot virus |
| A | QVX32675.1 | Tomato bushy stunt virus |
| B | QWC36209.1 | Pelargonium necrotic spot virus |
| B | QWC36204.1 | Pelargonium leaf curl virus |

Orange Cluster (Cluster No: 4)

| Set | Orange | Virus |
| --- | --- | --- |
| A | QWM97295.1 | Cymbidium mosaic virus |
| A | YP_010087744.1 | Euonymus yellow mottle associated virus |
| B | QXI69650.1 | Rhizoctonia solani dsRNA virus 18 |
| B | QXI69648.1 | Rhizoctonia solani dsRNA virus 14 |
| C | WP_216633687.1 | Oscillibacter sp. MSJ-2 |
| C | WP_215655026.1 | Pseudoflavonifractor sp. MCC625 |
| D | KAG8209403.1 | Trichophyton interdigitale |
| D | KAG7084114.1 | Colletotrichum scovillei |

Black Cluster (Cluster No: 5)

| **Set** | Black | Virus |
| --- | --- | --- |
| A | YP_010086016.1 | Tarumizu tick virus |
| A | YP_010086008.1 | Kundal virus |

Brown Cluster (Cluster No: 6)

| Set | Brown | Virus |
| --- | --- | --- |
| A | QXJ13539.1 | Mokola lyssavirus |
| A | QIK01206.1 | Rabies lyssavirus |
| A | YP_010086774.1 | Taiwan bat lyssavirus |

Teal Cluster (Cluster No: 7)

| Set | Teal | Virus |
| --- | --- | --- |
| Set A | QWT83775.1 | Odontoglossum ringspot virus |
| Set A | QWT83771.1 | Youcai mosaic virus |
| Set A | QWT83704.1 | Sunn-hemp mosaic virus |
| Set B | QXT50807.1 | Tomato brown rugose fruit virus |
| Set B | QWT83680.1 | Tobacco mosaic virus |
| Set B | QVX32625.1 | Tomato mottle mosaic virus |

Magneta Cluster (Cluster No: 8)

| **Set** | **Accession No.** | **Virus** |
| --- | --- | --- |
| Set A | YP_010086098.1 | Cacao virus |
| Set A | YP_010086191.1 | La Gloria virus |
| Set B | YP_010086093.1 | Itaporanga virus |
| Set B | YP_010086223.1 | Icoaraci virus |
| set C | QXP08778.1 | Wenzhou mammarenavirus |
| Set C | QVU02257.1 | Mammarenavirus AnRB3214 |

D) COBALT RESULTS

Green – A

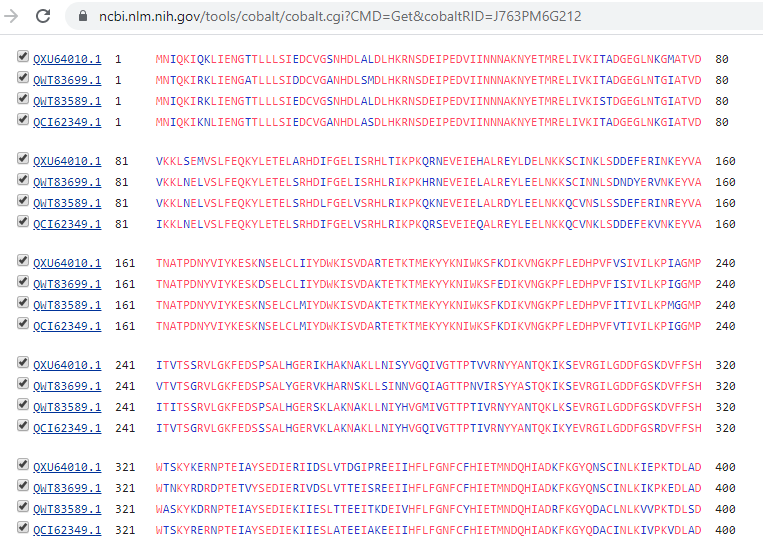

Green Set B:

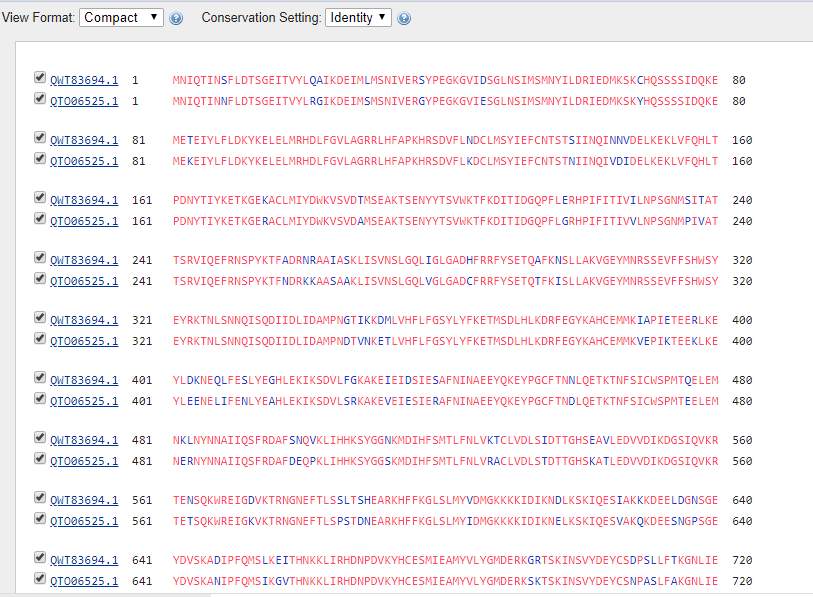

Green Set C:

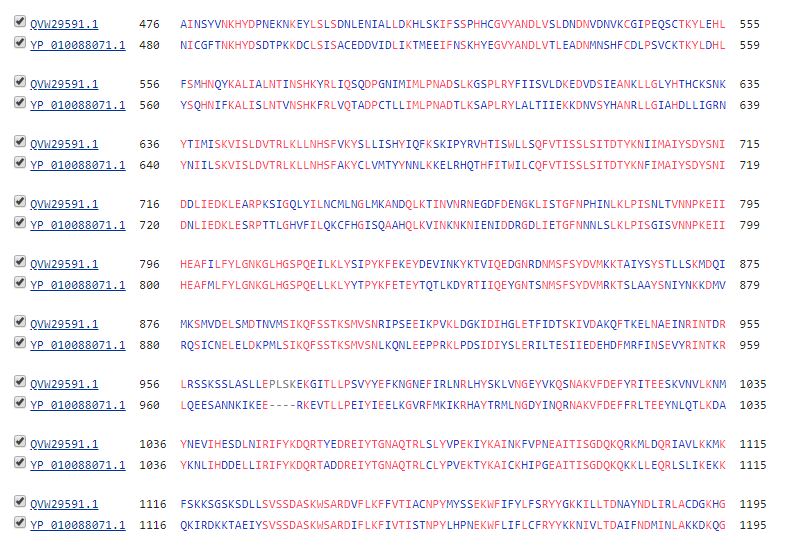

Red Cluster Set A:

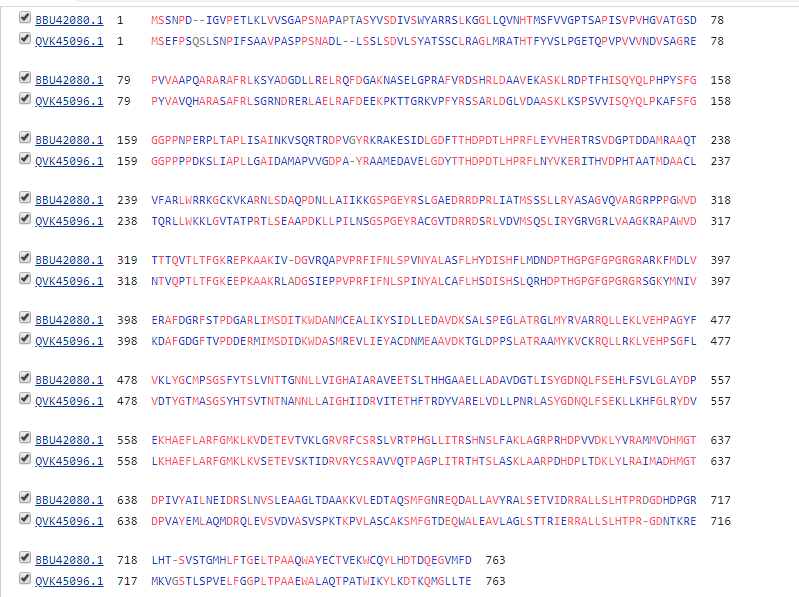

Red Cluster Set B:

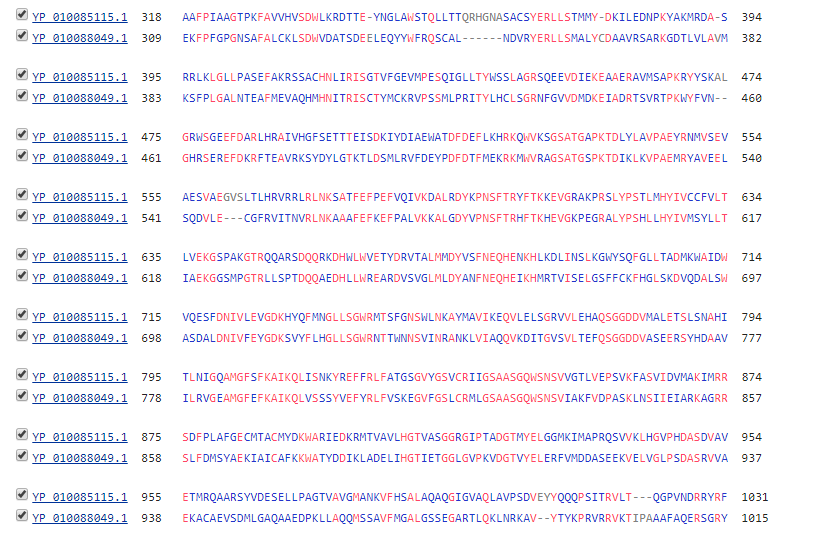

Blue set A:

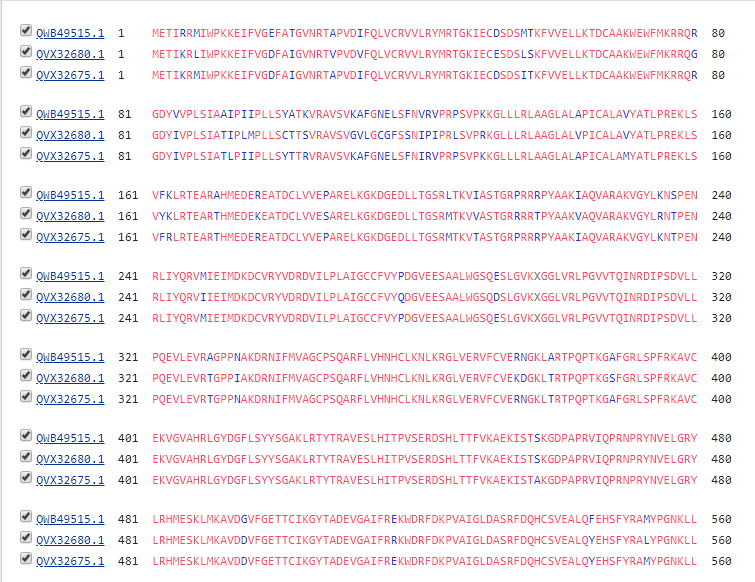

Blue Set B:

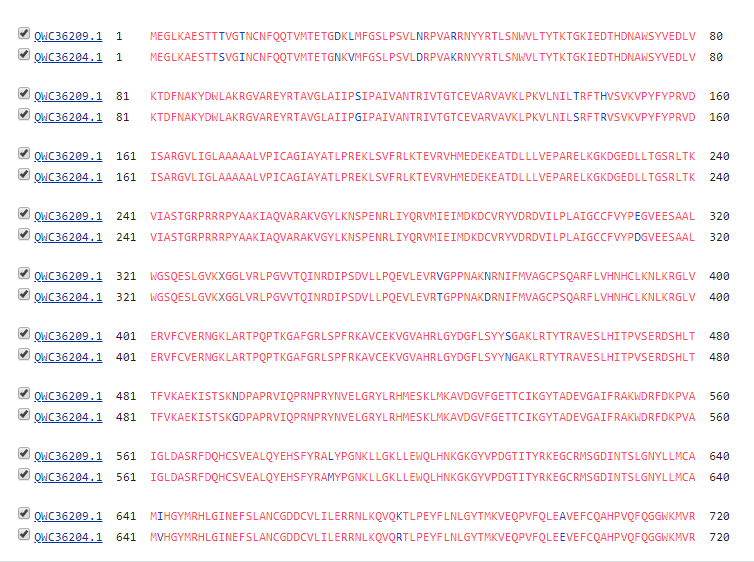

Orange Set A:

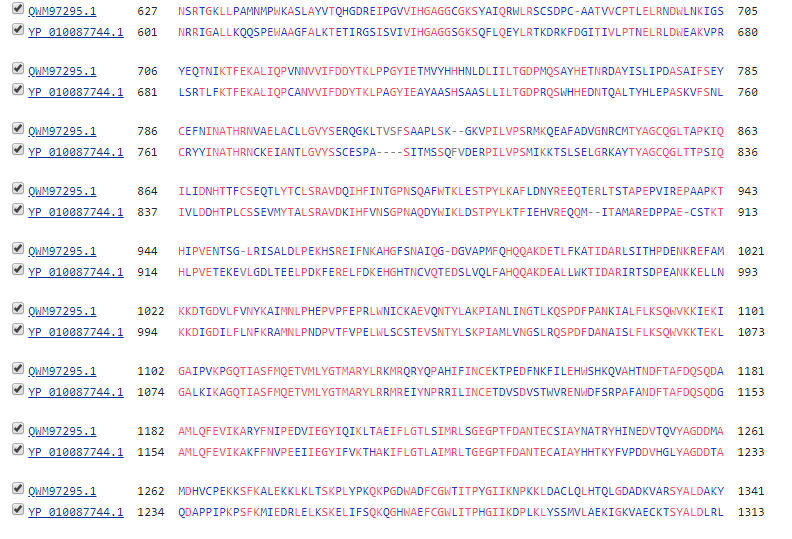

Orange Set B:

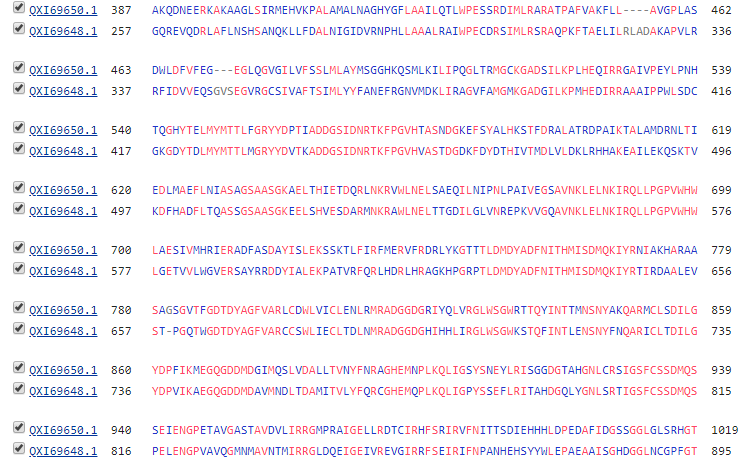

Orange Set C:

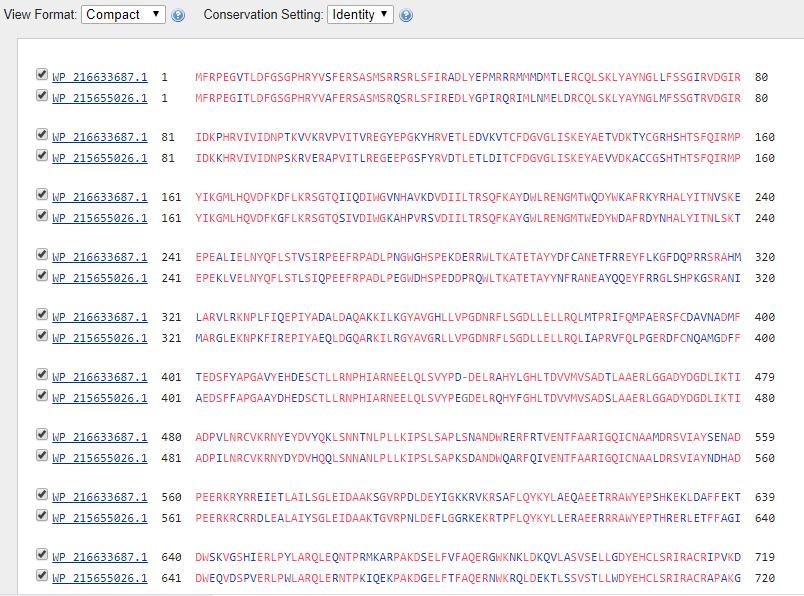

Orange Set D:

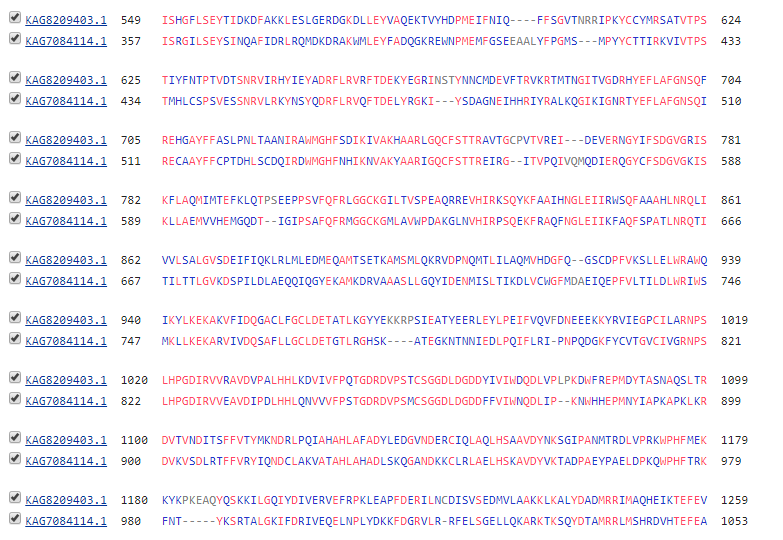

Black Set A:

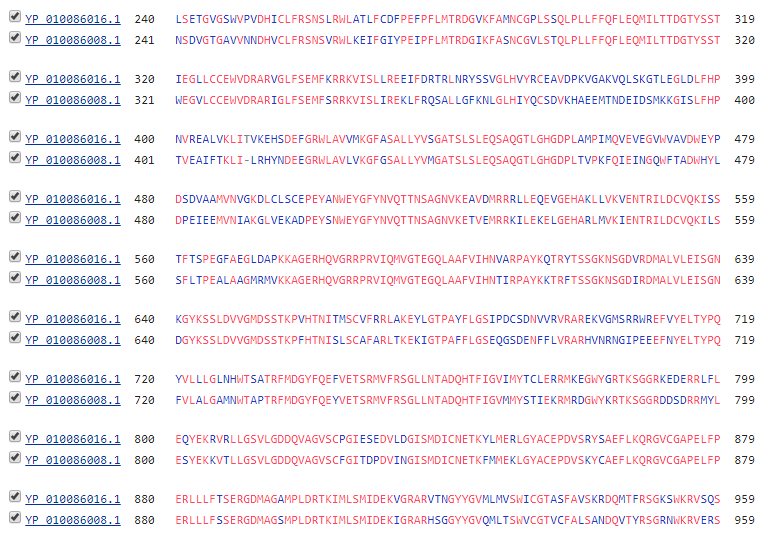

Brown Set A:

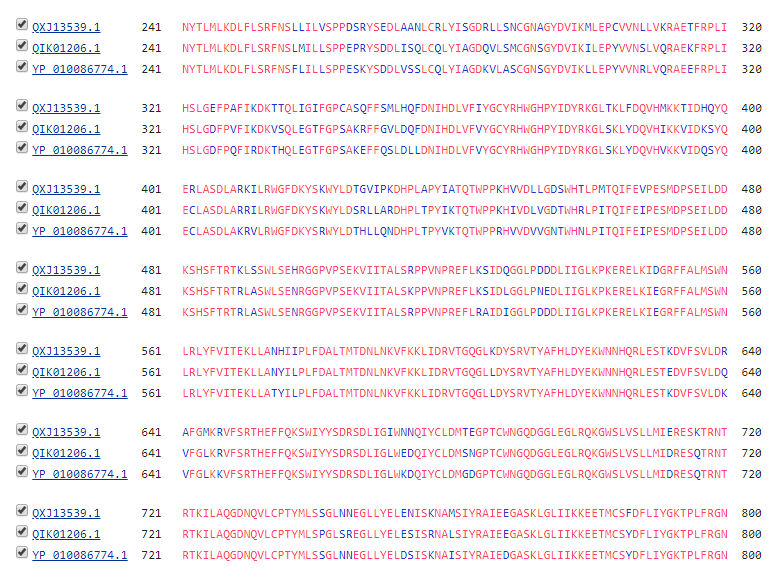

Teal set A:

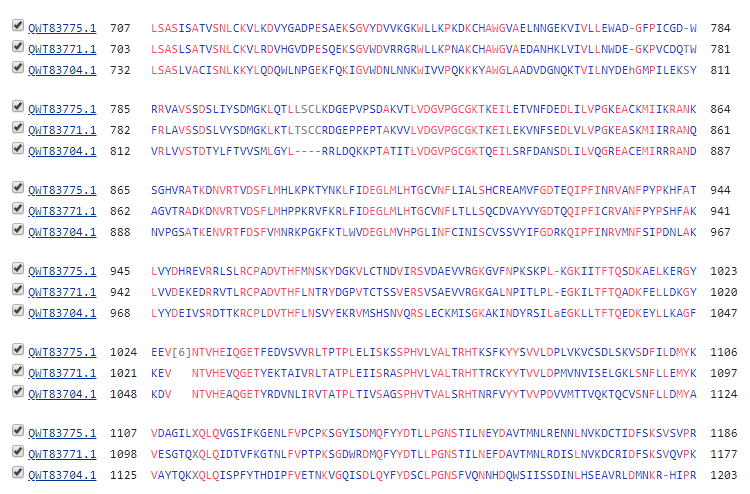

Teal Cluster B:

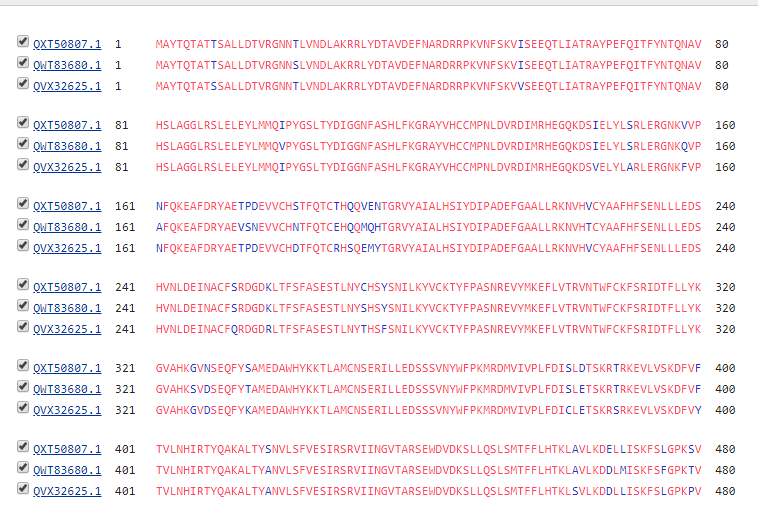

Magneta Set A:

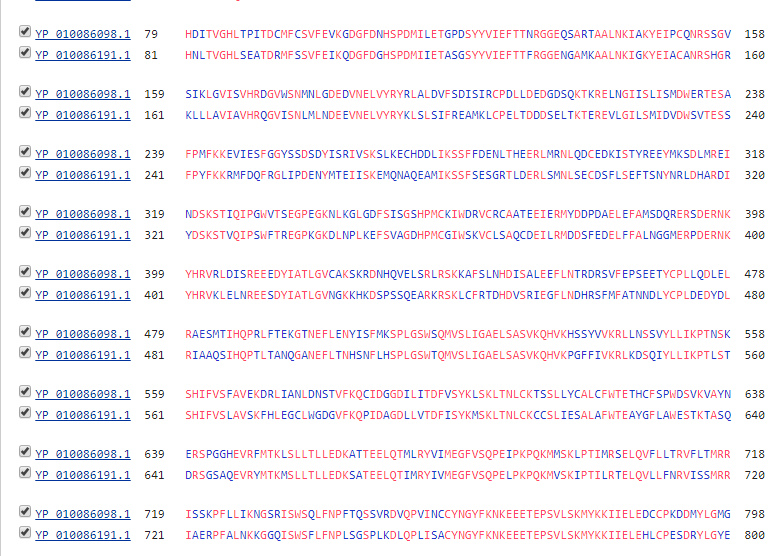

Magneta set B:

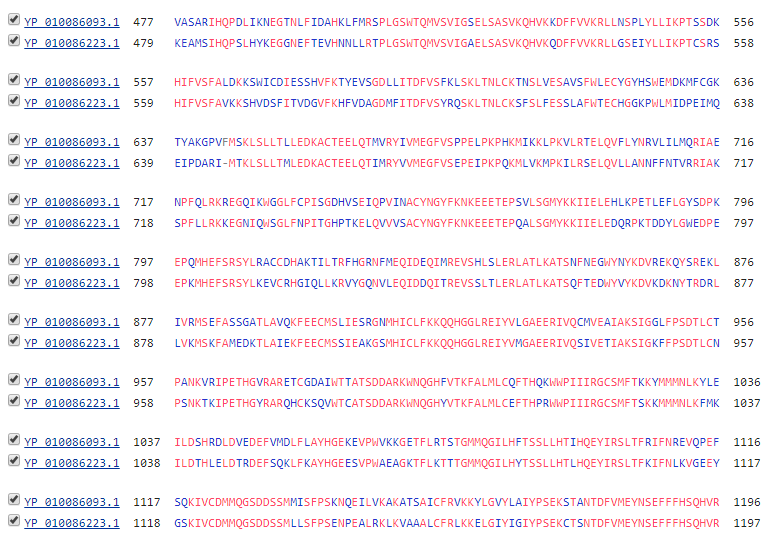

Magneta Set C:

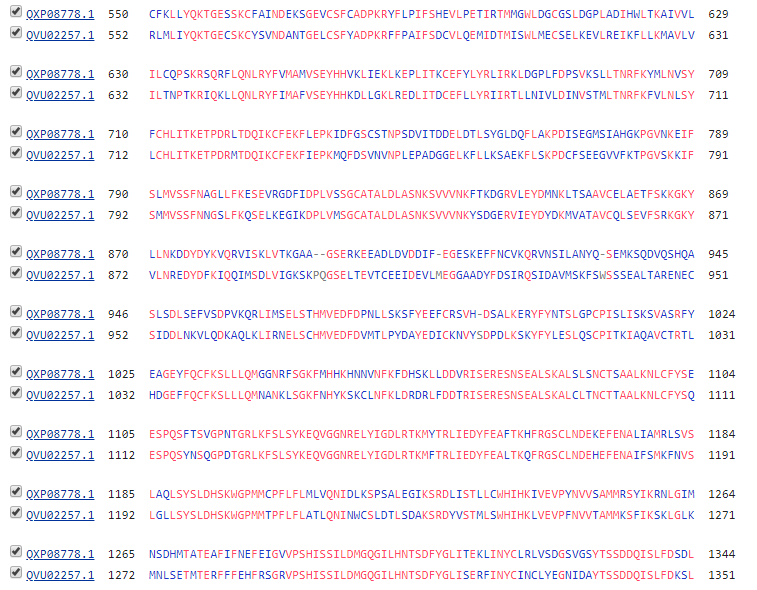

E) Codes:

Repository: Programs and Datasets : https://github.com/DSPavan/RdRP
